# Hypoxic gene expression in chronic hepatitis B infected patients is not observed in state-of-art *in vitro* and mouse infection models

**DOI:** 10.1101/2020.03.27.011841

**Authors:** PJ Liu, JM Harris, E Marchi, V D’Arienzo, T Michler, PAC Wing, A Magri, AM Ortega-Prieto, M van de Klundert, J Wettengel, D Durantel, M Dorner, P. Klenerman, U Protzer, S Giotis, JA McKeating

## Abstract

Hepatitis B virus (HBV) is the leading cause of hepatocellular carcinoma (HCC) worldwide. The prolyl hydroxylase domain (PHD)-hypoxia inducible factor (HIF) pathway is a key mammalian oxygen sensing pathway and is frequently perturbed by pathological states including infection and inflammation. We discovered a significant upregulation of hypoxia regulated gene transcripts in patients with chronic hepatitis B (CHB) in the absence of liver cirrhosis. We used state-of-the-art *in vitro* and *in vivo* HBV infection models to evaluate a role for HBV infection and the viral regulatory protein HBx to drive HIF-signalling. HBx had no significant impact on HIF expression or associated transcriptional activity under normoxic or hypoxic conditions. Furthermore, we found no evidence of hypoxia gene expression in HBV *de novo* infection, HBV infected human liver chimeric mice or transgenic mice with integrated HBV genome. Collectively, our data show clear evidence of hypoxia gene induction in CHB that is not recapitulated in existing models for acute HBV infection, suggesting a role for inflammatory mediators in promoting hypoxia gene expression.

## INTRODUCTION

HBV is a global health problem with more than 250 million people chronically infected and at least 780,000 deaths/year from HBV-related liver diseases such as liver cirrhosis and hepatocellular carcinoma (HCC)^1,2^. HBV replicates in hepatocytes within the liver and current anti-viral treatments suppress viral replication but are not curative, largely due to the persistence of the viral covalently closed circular DNA (cccDNA) reservoir^3^. Chronic hepatitis B (CHB) is a virus-associated, inflammatory liver disease and one of the leading causes of HCC^4^, one of the fastest rising and fourth most common cause of cancer related-death world-wide^5^. Curative therapies (tumour ablation, resection or liver transplantation) are dependent on early detection, however, the majority of HBV and non-viral associated HCC cases are diagnosed at a late stage often resulting in a poor prognosis^6^. Despite significant advances in our understanding of the HBV replicative life cycle, the mechanisms underlying HCC pathogenesis are not well defined^7^.

Although liver cirrhosis is a major risk factor for developing HCC, however, 10-20% of HBV infected patients that develop HCC are non-cirrhotic, highlighting a role for HBV to promote carcinogenesis via direct and indirect inflammatory mechanisms^7^. Three major and non-exclusive viral-dependent pathways have been proposed: (i) integration of viral DNA into the host genome; (ii) expression of viral oncogenic proteins and (iii) viral-driven changes in host gene transcription (reviewed in^8^). The viral encoded regulatory hepatitis B X protein (HBx) has been reported to promote the expression of both viral and selected host genes, where a recent study reported HBx binding to >5,000 host genes with diverse roles in metabolism, chromatin maintenance and carcinogenesis^9^. There is clearly an urgent need to increase our understanding of HBV mediated carcinogenesis to support the development of tools to identify CHB patients at risk of HCC development.

The liver receives oxygenated blood from the hepatic artery and oxygen-depleted blood via the hepatic portal vein, resulting in an oxygen gradient of 8-4% across the periportal and pericentral areas, respectively^10^. This oxygen gradient has been reported to associate with liver zonation, a phenomenon where hepatocytes show distinct functional and structural heterogeneity across the parenchyma^11,12^. Recent single-cell RNA sequencing analysis of the mouse liver highlight a major role for hypoxic and Wnt signalling pathways to shape liver zonation profiles in the normal healthy liver with an enrichment of hypoxic gene expression in the pericentral area^13^. Importantly, this oxygen gradient is readily perturbed in pathological states such as infection, inflammation and cirrhosis^14^. One of the most well studied oxygen sensing mechanisms is the hypoxia inducible factor (HIF) pathway^15^. As HIF-signalling pathways are altered in many diseases, including cancer and inflammatory conditions, pharmacological approaches to modulate HIF activity offer promising therapeutic opportunities^16,17^. When oxygen is abundant, newly synthesised HIFα subunits, including HIF-1α and HIF-2α isomers, are rapidly hydroxylated by prolyl-hydroxylase domain (PHD) proteins and targeted for poly-ubiquitination and proteasomal degradation. In contrast when oxygen is limited these HIFα subunits translocate to the nucleus, dimerize with HIF-β and positively regulate the transcription of a myriad of host genes involved in cell metabolism, proliferation, angiogenesis and immune regulation. Dai *et al.* reported that increased HIF-1α mRNA and protein expression in HCC are prognostic for more advanced disease stages and poor overall survival post-surgical tumour resection^18^. Furthermore, Xiang *et al.* and Zheng *et al.* showed that HIF-1α protein expression is predictive of HCC lymph node metastasis and vascular invasion^19,20^. Thus, HIF signalling could have an important role in progressive liver disease and HCC development^14^.

In addition to hypoxia, inflammation, oxidative stress and viral infection can promote HIF-transcriptional activity. The host inflammatory mediators nuclear factor-κB (NF-κB) and tumor necrosis factor-α (TNF-α) induce HIF-1α transcription^21,22^. Reactive oxygen species (ROS) produced by inflammatory cells provide a further mechanism for inflammation-driven HIF-signalling^23-25^. Several viruses induce the HIF signaling pathway including hepatitis C virus^26-28^, human papillomavirus^29^, Kaposi sarcoma-associated herpesvirus^30^ and human cytomegalovirus^31^. Several reports have suggested that HBx can interact with and stabilize HIFs^32-40^, however, this proposed HBx-HIF interplay awaits validation in HBV replication *in vitro* and *in vivo* model systems.

In this study, we report a significant upregulation of hypoxic gene expression in a cohort of chronic HBV infected patients^41^. However, our studies to investigate the underlying mechanism using state-of-the-art *in vitro* and *in vivo* HBV transgenic mice and human liver chimeric mice models show limited evidence of hypoxic gene expression. These studies highlight a major role of liver inflammation and a complex interplay between HBV and HIF signalling in the chronic infected liver that is not recapitulated by current infection-competent model systems. Collectively, our data show clear evidence of hypoxia-driven gene expression in CHB in the absence of cirrhosis or HCC development that may play a role in driving hepatocarcinogenesis.

## RESULTS

### Increased hypoxia gene signature in chronic hepatitis B

To determine whether there is any evidence of hypoxic associated transcription in CHB we used a published RNA-seq transcriptome of hypoxic HepG2 cells^42^ (cultured at 0.5% oxygen for 16 hours) to identify 80 genes with a greater than 2-fold increase in transcript levels (false discovery rate (FDR) of < 0.05) (**Supplementary Table 1**). We assessed the expression profile of these hypoxic regulated genes using a published liver transcriptome (Affymetrix-microarray) data set obtained from a cohort of chronic HBV infected patients (CHB, n=90) that were free of cirrhosis or HCC and uninfected control subjects (n=6 healthy)^41^. We excluded any patients that had no detectable serum HBV DNA. We noted an increase in HIF-1α mRNA levels in the CHB patients compared to control subjects (Log2 FC = 2.648, p=0.005). Gene set enrichment analysis (GSEA) showed a significant enrichment in the hypoxia gene set in the infected patients (**Fig.1a**). To extend this observation we used GSEA to screen the CHB cohort for expression of 67 HIF and hypoxia signatures obtained from the Molecular Signatures Database (MSigDB v 7.0, https://www.gsea-msigdb.org/gsea/msigdb/). We observed an enrichment of over 50% of these gene sets in CHB cohort (FDR<0.25), confirming increased expression of hypoxic genes in CHB liver (**Fig.1b**). To evaluate other enriched pathways in CHB liver, GSEA was carried out using the Hallmark gene sets from MSigDB. This data base is curated to have minimal overlap between categories, reducing noise and redundancy and summarize specific cell states or biological processes. This analysis identified genes associated with allograft rejection as the most significantly upregulated gene set in CHB. Interestingly HIF-1α was one of the leading-edge genes in this subset; contributing significantly to the core enrichment score. Moreover, we noted a significant increase in inflammatory signaling pathways in CHB liver: ‘TNFA signaling via NF-kB’, ‘Inflammatory Response’ and ‘Interferon Gamma Response’ (**Fig.1c**). In summary, these data show increased hypoxic gene signatures in CHB liver that associates with an activation of inflammatory pathways.

**Fig. 1.**
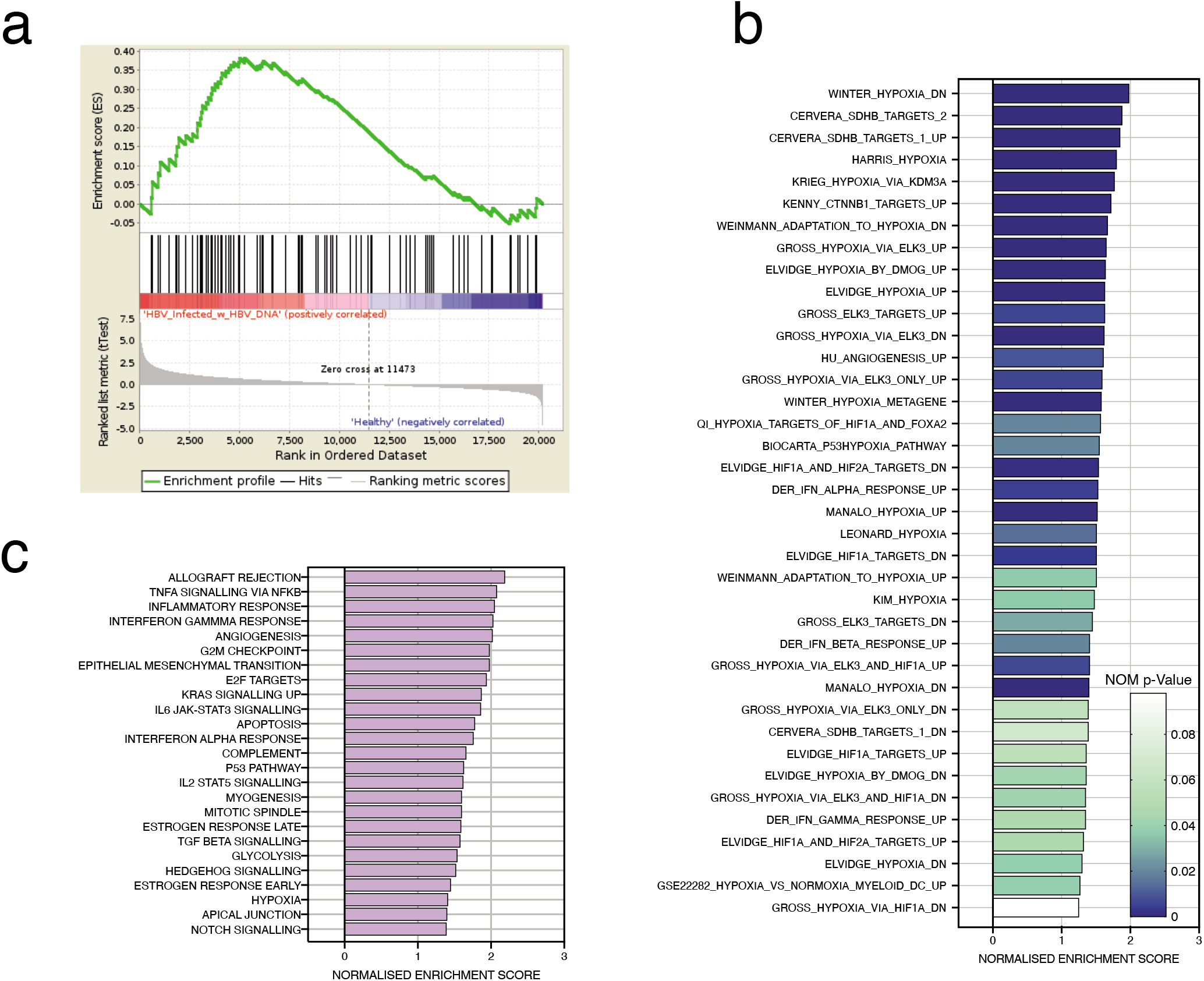
Increased hypoxia gene expression in CHB. GSEA shows a significant enrichment of HepG2 defined hypoxic genes^42^ in chronic HBV infected patients^41^ vs healthy patients (FDR = 0.06). The gene set was based on Fold Change > 2, and FDR < 0.05; 80 genes satisfied these criteria and are listed in supplementary Table 1 (**a**). HIF and hypoxia gene signatures from Molecular Signatures Database (MSigDB v 7.0, https://www.gsea-msigdb.org/gsea/msigdb/) were assessed in the CHB transcriptomic data set and 38 significantly upregulated genes identified (FDR<0.25) in CHB patient subsets ranked by Net Enrichment Score (NES) (**b**). Using the upregulated MSigDB hallmark gene sets, GSEA was used to identify the most upregulated pathways in the CHB cohort. 33 gene sets were significantly enriched (FDR<0.25). The image shows the top 25 most significantly enriched gene sets, ranked by NES (**c**).

### Limited evidence for HBx to stabilise HIF-1α or HIF-2α expression or associated transcriptional activity *in vitro*

As HBx is the major viral encoded transcriptional activator we wanted to assess its role in stabilizing HIFs and used the bipotent HepaRG cell line engineered to express HBx (HepaRG-HBxWT) under a tetracycline (Tet) inducible promoter^43,44^ and confirmed HBx expression (**Fig.2a**). HBx in this model system is functionally active and can restore the replication of mutant viruses lacking HBx^44^. HBx promotes viral transcription by degrading the host structural maintenance of chromosomes (Smc) complex Smc5/6^45^ and we confirmed the loss of Smc6 expression in Tet-induced HepaRG-HBxWT cells (**Fig.2a**). As a control for these experiments we generated HepaRG cells encoding HBx with three nonsense mutations (HepaRG-HBx_STOP_). To assess whether HBx can promote or stabilize HIF expression we treated HepaRG-HBxWT or HepaRG-HBx_STOP_ cells with Tet and cultured at 1% oxygen, a typical oxygen concentration used to model hypoxia ex vivo, or standard ‘normoxic’ laboratory conditions of 20% oxygen for 24h. HBx had minimal impact on HIF-1α or HIF-2α protein (**Fig.2b**) or mRNA levels (**Fig.2c**) in HepaRG cells cultured at 20% oxygen. Culturing HepaRG-HBxWT or HepaRG-HBx_STOP_ cells under 1% oxygen confirmed HIF-1α or HIF-2α expression and importantly showed a negligible effect of HBx on either HIF isoform (**Fig.2b**). To assess whether HBx altered HIF transcriptional activity we quantified the mRNA levels of four HIF-regulated host genes (*CAIX, BNIP3, VEGFA* or *GLUT1*) (**Fig.2c**) and CAIX protein expression (**Fig.2b**) and observed no differences. Under normoxic conditions HIFs are hydroxylated by the oxygen-dependent PHDs and targeted for proteosomal degradation. Oxygen reperfusion of hypoxic cells results in a time-dependent loss of HIFs and we assessed whether the presence of HBx could alter the kinetics of HIF expression. A comparable decrease in HIF-1α and HIF-2α proteins was seen after 10-20 mins of oxygen reperfusion in both Tet treated and untreated cells (**Fig.2d**), demonstrating that HBx has a negligible effect on the kinetics of HIF degradation. In summary, we demonstrate that HepaRG cells are responsive to low oxygen and show a significant increase in hypoxia-associated gene transcription, this effect was not impacted by the co-expression of HBx.

**Fig. 2.**
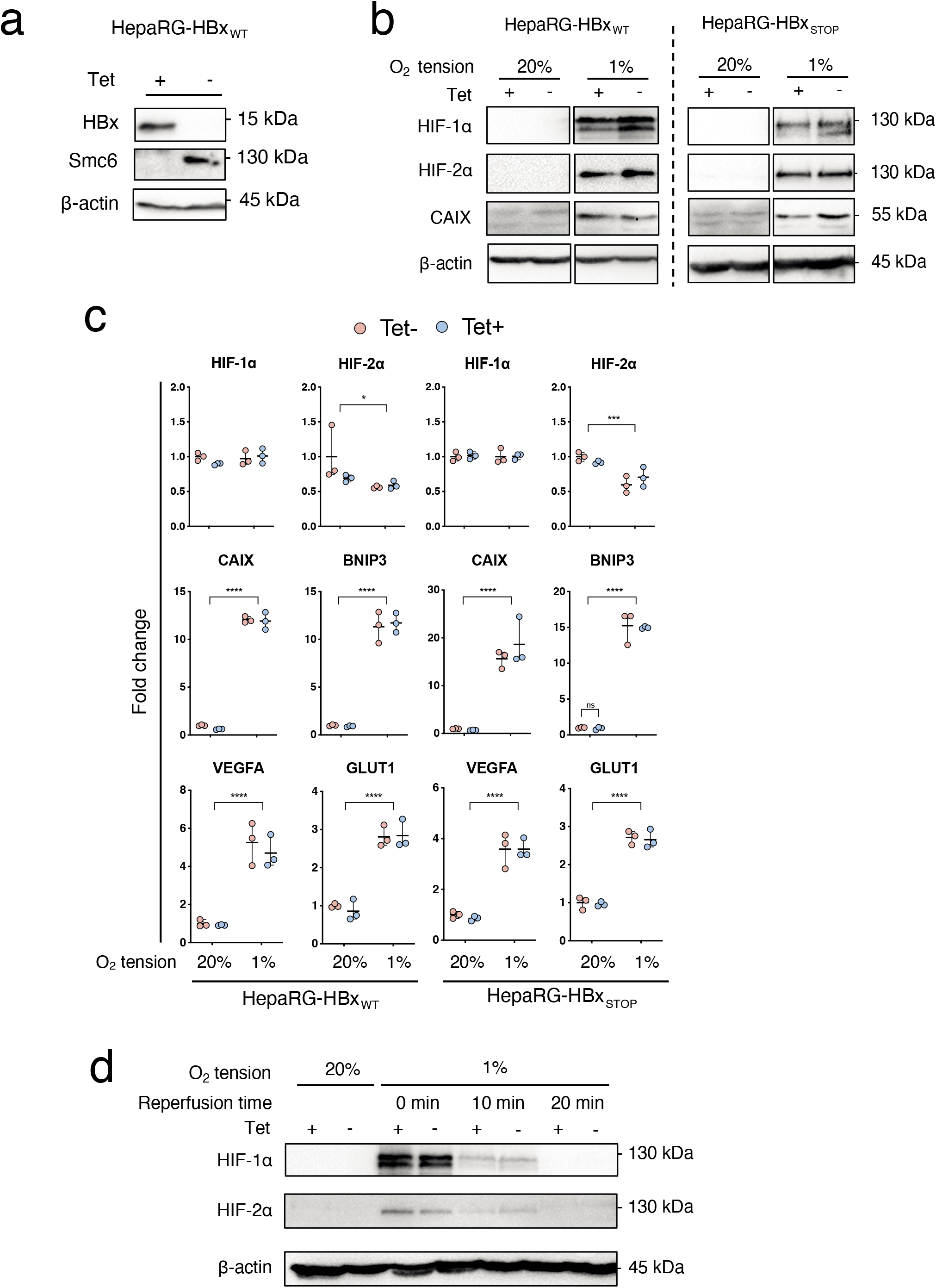
Effect of HBx on HIF expression and transcriptional activity in HepaRG cells. HepaRG cells encoding HBx were incubated with Tet (50 μM) for 24 h and HBx protein and Smc6 expression detected by western blot (**a**). HepaRG cells encoding WT or mutated HBx (STOP) were incubated with or without Tet (50 μM, 24 h) and cultured under 20% or 1% oxygen conditions for 24h. Cells were lysed and expression of HIF-1, HIF-2, Carbonic anhydrase IX (*CAIX*) and housekeeping gene B-actin assessed by western blotting (**b**) and mRNA levels of HIF-1, HIF-2 and several HIF target genes (*CAIX, BNIP3, VEGFA* and *GLUT1*) quantified by qPCR (**c**). HepaRG cells encoding wild type HBx were incubated with Tet (50 μM, 24 h) and cultured at 20% or 1% oxygen for 24h. The hypoxic cultures were returned to 20% oxygen. After 10 or 20 mins, cells were lysed and screened for HIF-1α or HIF-1α and housekeeping gene B-actin expression (**d**). The data is shown from a single experiment and is representative of three independent experiments and represents mean ! standard deviation. Normality distribution was assessed by D’Agostino-Pearson test; 2-way-ANOVA with Bonferroni correction was applied with p<0.05 deemed as significant.

To further investigate a role for HBx to stabilize HIF-1α we used an adenoviral vector engineered to express HBx (Ad-HBx) and confirmed HBx expression and Smc6 degradation. Additionally, transduction of HepG2-NTCP cells with Ad-HBx restored the replication of an HBV virus with a mutated HBx open reading frame (HBV_X-_) further demonstrating its functional activity (**Fig.3a**). HepG2-NTCP cells transduced with Ad-HBx or Ad-OVA (adenoviral vector expressing ovalbumin) were cultured at 20% or 1% oxygen and cells harvested over a 48h period. We confirmed HBx expression 24h post-transduction (**Fig.3b**) and observed expression of HIF-1α after 8h at 1% O2. Comparable expression levels of HIF-1α were noted in both Ad-HBx and Ad-OVA transduced cells, demonstrating a negligible effect of HBx on HIF-1α induction. These results further highlight a minimal role of HBx in regulating HIF1α or HIF2α mRNA or protein expression.

**Fig. 3.**
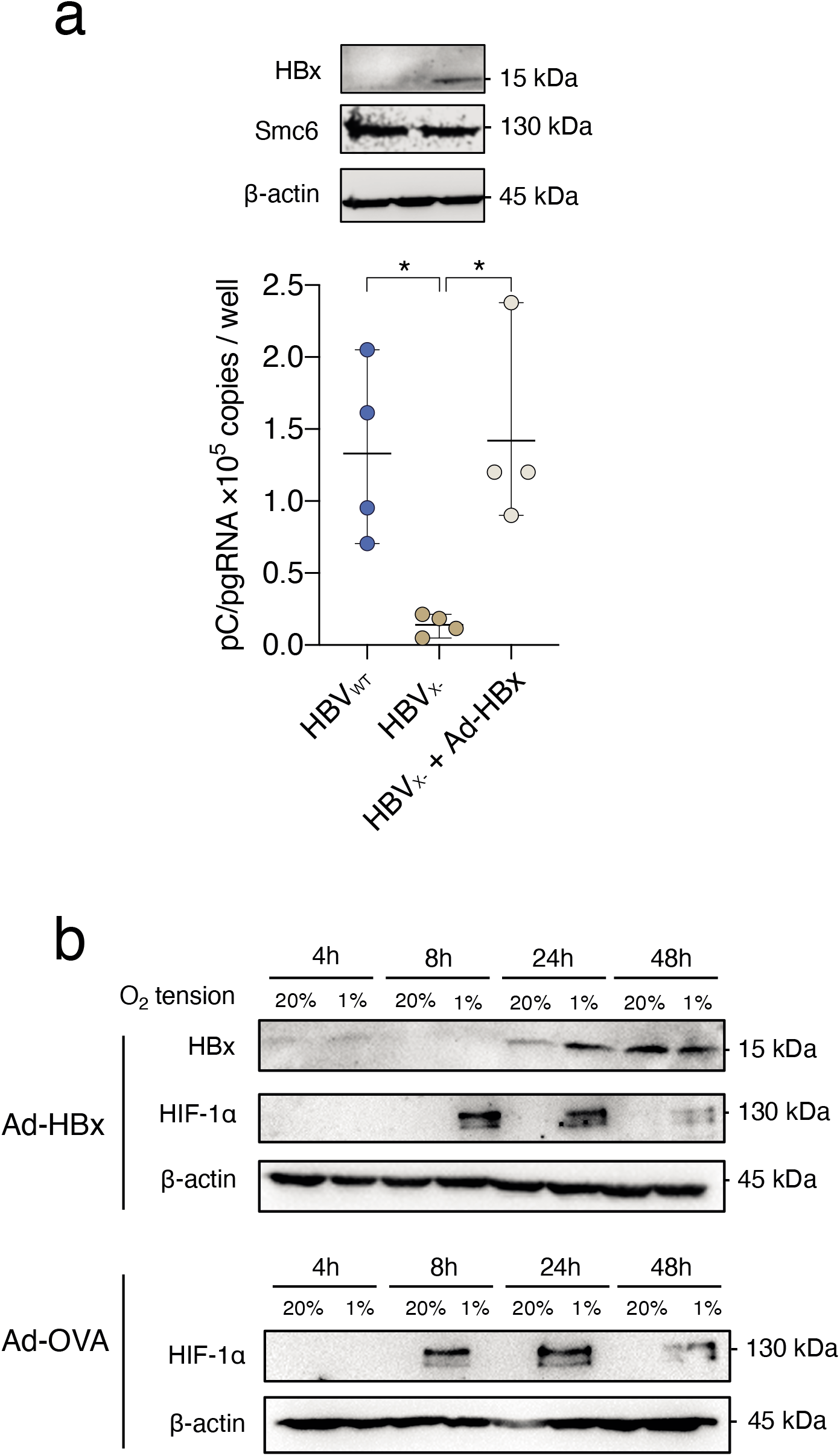
Effect of HBx expression on HIF expression and transcriptional activity in HepG2 cells. HepG2-NTCP cells were transduced with Ad-HBx and 24h later the cells were probed for HBx and Smc6 expression. In parallel experiments HepG2-NTCP cells were infected with HBV or a mutated virus lacking HBx (HBV_X-_) (MOI of 200) in the presence or absence of Ad-HBx and the major viral transcript, pregenomic RNA measured at 6 days post-infection (**a**). HepG2-NTCP cells were transduced with Ad-HBx or Ad-OVA and HBx and HIF-1 expression assessed at selected times after culturing at either 20% or 1% oxygen (**b**). Data is shown from a single experiment and is representative of three independent experiments and mean data is presented. Normality distribution was assessed by D’Agostino-Pearson test; 2-way-ANOVA with Bonferroni correction was applied with p<0.05 deemed as significant.

### Studying HIF transcriptional activity in HBV transgenic mice

Since HBV can only infect humans and hominoid primates, no immune competent animal models are available that support natural HBV infection. One of the most-widely used murine models for studying CHB are transgenic mice expressing HBV from a single integrated genome (HBVtg). HBVtg mice have been reported to develop HCC that show similar chromosomal aberrations and gene expression patterns to human HBV-associated HCC^46^. To study the effect of HBV on HIF transcriptional activity in this model system, HBVtg mice were treated with lipid nanoparticle complexed, liver-targeted siRNAs designed to silence all HBV transcripts (siHBV)^47^ or with an unspecific control siRNA (siCtrl). The HBV-specific siRNA led to effective HBV silencing with greater than 95% reduction in HBeAg in the serum (**Fig.4a**) and viral transcripts in the liver (**Fig.4b**). However, silencing HBV mRNAs and antigens had no impact on HIF regulated gene transcripts (*CAIX, VEGFA, GLUT1* and *PHD2*) (**Fig.4c**). These studies suggest a minimal role of HBV encoded proteins or RNAs in promoting HIF transcriptional activity.

**Fig. 4.**
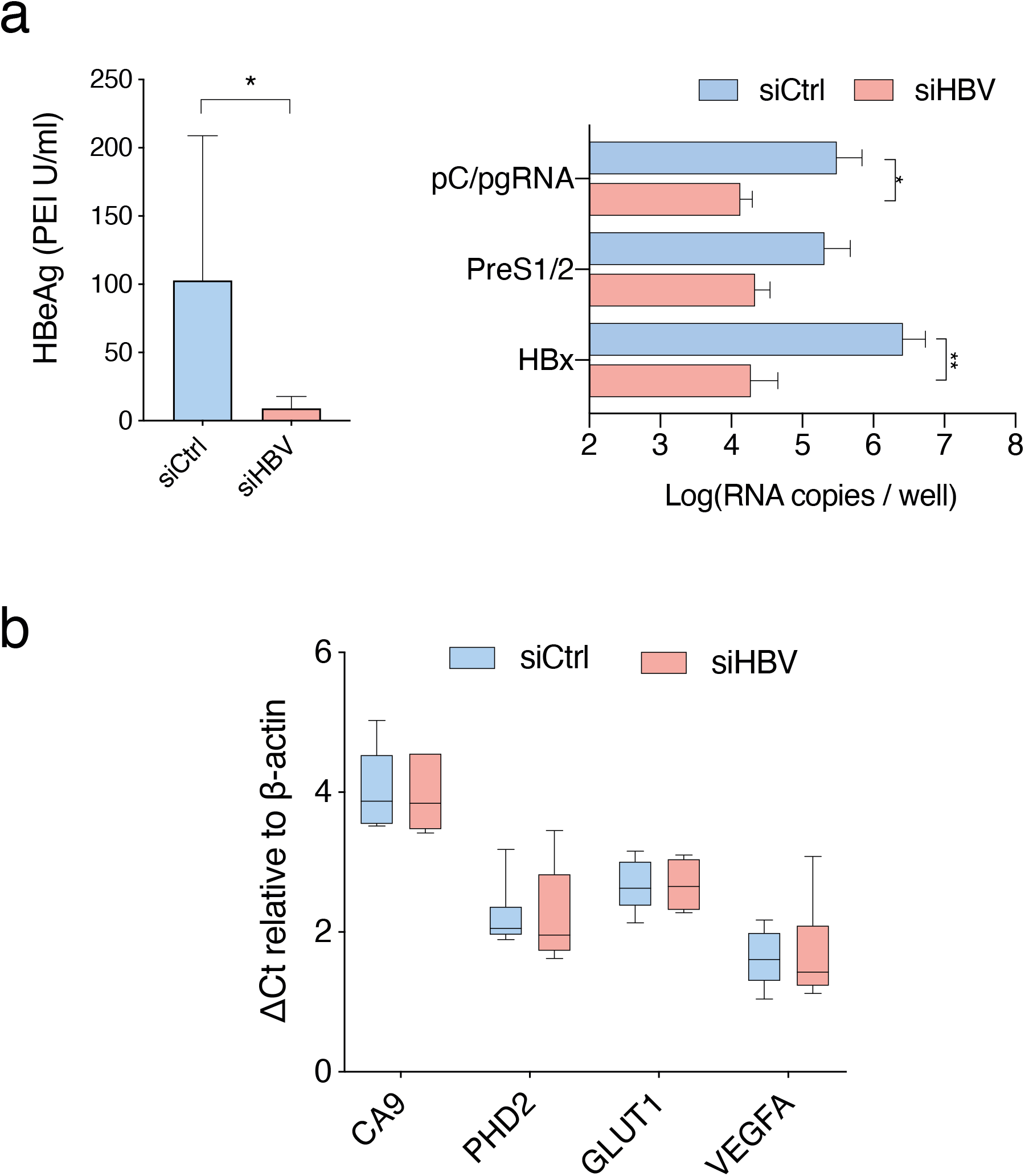
Effect of silencing viral transcription in HBV transgenic mice on hypoxia target gene transcripts. HBV transgenic mice (n=6 per group) were treated with liver directed siRNAs targeting the HBx region (siHBV) which is commonly shared by all viral RNAs or with a control siRNA (siCtrl). Seven days later we assessed the efficacy of siHBV silencing by quantifying: serum HBeAg levels (**a**), HBV RNAs in the liver (**b**) and hypoxia target gene *(CAIX, VEGFA, GLUT1* and *PHD2)* RNAs (**d**). Hypoxia target genes values are expressed as ΔCt values by subtracting the Ct value of the housekeeping gene ß-actin from Ct value of the gene of interest. Mann Whitney test (a) or 2-way-ANOVA with Bonferroni correction (b) test were applied with p<0.05 deemed as significant.

### Studying HIF transcriptional activity in HBV infected hepatocytes and human liver chimeric mice

To complement the HBx studies described above we investigated the effect of HBV infection on HIF oxygen sensing pathways in current state-of-the-art *in vitro* and *in vivo* models. HepG2-NTCP cells were infected with HBV and cultured under normoxic conditions and sampled after 3 and 9 days to assess HIF-1α or HIF-2α expression. HBV gene expression was confirmed by measuring HBeAg (53.96 ± 2.7 IU/mL) and HBsAg (12.63 ± 4.4 IU/mL), however, we failed to detect either HIF or CAIX expression in the infected or non-infected cells (**Fig.5a**). As a control we treated HepG2-NTCP cells with a HIF PHD inhibitor (FG4592 at 30 μM) and demonstrated HIF protein expression (**Fig.5a**). Analyzing a published transcriptomic RNA-seq data from HBV infected primary human hepatocytes^48^ showed no evidence of hypoxic gene upregulation (**Fig.5b**). To further validate our conclusions we used the chimeric human liver FNRG mouse model^49^ to assess whether HBV infection would induce HIF signaling in this model. Female FNRG mice^49^ between 8-12 weeks of age were transplanted with 0.5×10^6^ cryopreserved adult human hepatocytes by intrasplenic injection and monitored for engraftment by measuring human albumin levels in the serum (at least 0.1mg human albumin per mL in peripheral blood). Engrafted animals were infected with 0.5 million genome equivalent (GE) copies of HBV per mouse and were monitored for HBV replication. Once stable viremia was established (minimum 5×10^7^ GE mL^−1^ of serum) the mice were sacrificed and livers harvested from HBV infected (n=4) and uninfected (n=3) animals for RNA isolation and RNA-sequencing. Analysing these RNA-seq data sets showed minimal evidence for an increase in hypoxic transcriptional activity in the HBV infected livers (**Fig.5b**). For comparative purposes, we show that hypoxic genes were upregulated in the CHB cohort^41^ (**Fig.5b**), demonstrating the influence of inflammation on gene regulation and highlighting the limitations of current HBV replication models to model CHB.

**Fig. 5.**
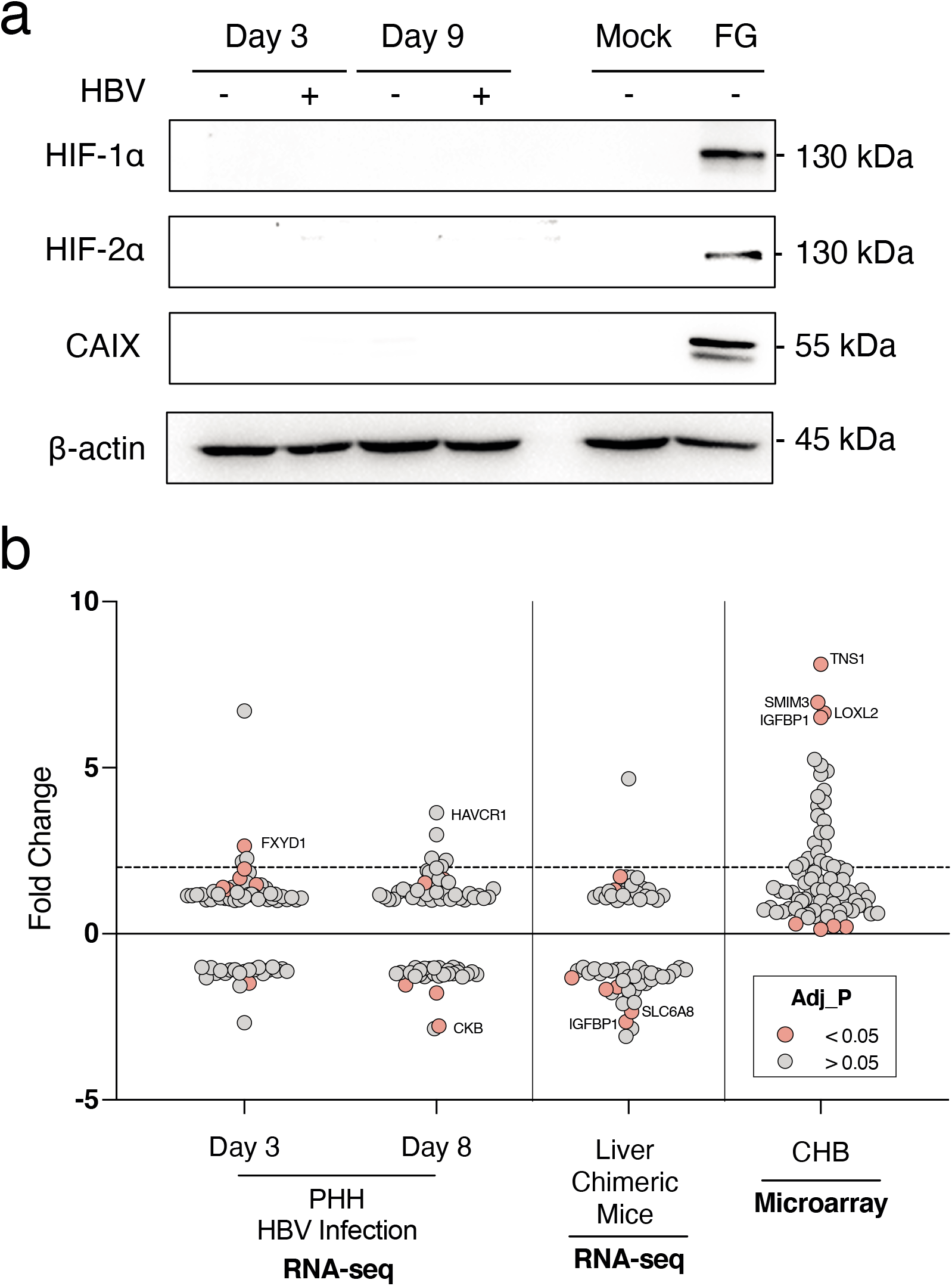
Comparing hypoxia gene signatures in HBV infected hepatocytes and humanized liver chimeric mice. Mock or HBV-infected HepG2-NTCP cells (MOI 200) were harvested after 3 or 9 days, lysed and assessed for HIF-1α, HIF-2α or CAIX expression and the housekeeping gene B-actin by western blotting. As a positive control HepG2-NTCP cells were treated with the HIF PHD inhibitor FG4592 (FG, 30 μM) for 24h and protein lysates analysed by western blotting (**a**). Induction of hypoxic genes (Supplementary Tabe 1) in transcriptomic data of HBV infected primary human hepatocytes^48^, HBV infected human liver chimeric mice and a CHB cohort (**b**). Fold change was calculated for each of the 80 genes in HBV infection against the healthy controls, where the dotted line represents a 2 fold change. For the CHB cohort, fold change was calculated from the raw Affymetrix, differential expression was tested using multiple t-tests and significance determined by p value <0.05.

## DISCUSSION

In this study we identified increased hypoxia gene signatures in a CHB cohort in the absence of cirrhosis or HCC. We noted an increase in HIF-1α mRNA levels, consistent with their transcriptional regulation by inflammatory mediators such as TNFα. Given previous reports that HBx can stabilize HIFs^32-40,50^, we investigated whether functionally active HBx could regulate endogenous HIF-1α and HIF-2α mRNA, protein and transcriptional activity *in vitro*. We found minimal evidence for HBx regulation of HIFs in three independent model systems: an inducible HepaRG-HBx cell line; an Ad-HBx transduced cell, and in *de novo* infection of HepG2-NTCP or PHHs. To reconcile our observations with previous publications it is relevant to recognize the differences from earlier studies. Firstly, due to the technical difficulties in visualizing HBx by western blotting or immunofluorescent imaging, many of the earlier studies did not confirm HBx expression. Secondly, the majority of studies did not validate the functional activity of the expressed HBx protein. Finally, several studies assessed HBx stabilization of HIF-1α using transient plasmid transfection systems with hypoxia reporter constructs, rather than directly measuring HIF expression and HIF target gene modulation. Given our current knowledge that HBx degrades Smc6 that silences episomal DNA transcription, the interpretation of these earlier plasmid based systems^51^ is now uncertain. Since we have directly confirmed expression and function of HBx in our *in vitro* models and quantified endogenous HIF transcriptional activity under normoxic or hypoxic condtions we are confident that HBx does not modulate HIF expression or transcriptional activity in the model systems used.

Guerrieri et al. 2017 identified and validated a role for HBx in regulating genes involved in endocytosis, predominantly members of the Ras-related in brain (Rab) family^9^. Anti-HBx chromatin immunoprecipitation studies identified HBx binding sites that included RAB1A, RAB2B and RAB5B promotors and none of the validated Rab genes were listed in our hypoxic gene set (**Supplementary Fig.1**). Furthermore, GSEA of the CHB cohort or screening reactome gene sets showed only a modest enrichment in the ‘Transferrin Endocytosis’ pathway (**Fig.1c**), suggesting a minimal overlap between HBx and HIF regulated genes.

Our results support a model where HBV infection associated inflammatory responses promote HIF expression and these complex virus-cell interactions are not recapitulated by simple *in vitro* culture systems, HBV transgenic mice or immunodeficient SCID human liver chimeric mouse models. Our bio-informatic analysis identified 25 hypoxia upregulated genes in chronic HBV infected patients, including *LOXL2, SMIM3, TNS1*, and *IGFBP1*. Notably, *LOXL2* overexpression in HCC was previously associated with high tumour grade, metastasis, and poor patient overall and disease-free survival^52^. LOXL2 was shown to mediate its pathogenic effects in HCC angiogenesis via vasculogenic mimicry signalling, cytoskeleton reorganization, and bone-marrow derived cell recruitment^52,53^. In fact, hypoxia and HIF-1α signalling have been identified as key regulators of LOXL2 and driver of its pathogenesis, consistent with our observations^53,54^. Another significantly upregulated gene in chronic HBV patients, *IGFBP1*, was recently reported to be a HIF-2α regulated gene *in vitro* and *in vivo* model systems^55^. Furthermore, *IGFBP1* is a known NFκB target gene and is induced by HBV infection^56^. These data suggest co-regulation of *IGFBP1* by inflammatory pathways including NFκB and oxygen sensing mechanisms such as HIF signalling, which is consistent with our observation of inflammatory gene enrichment associating with hypoxia gene signature in CHB.

Our observation of increased hypoxic gene signature expression in CHB patients offers an important insight on HBV disease stage stratification and suggest areas for bio-marker discovery for early HCC detection. This is in agreement with previous studies that have associated higher HIFα mRNA and protein expression in HCC with worse prognostic outcomes for HCC patients ^18-20^. Moreover, as the liver is a naturally physiologically low oxygen environment, future investigations exploring how oxygen sensing pathways regulate HBV replication and pathogenesis may identify novel therapeutic targets

## MATERIALS AND METHODS

### Cell lines and reagents

HepaRG cells expressing HBx under the control of a Tetracycline inducible promoter were cultured in Williams E medium supplemented with 10% FBS, 50 U penicillin/streptomycin mL^−1^, 5 μg human insulin mL^−1^ and 5×10^−7^ M hydrocortisone hemisuccinate (Sigma). As a control we generated HepaRG cells expressing an inactive HBx null mutant (HepaRG-HBx_STOP_) where three point nonsense mutations (relative to EcoRI site: C to A, 1393nt; C to A, 1396nt and C to T, 1397nt) were introduced to generate three stop codons (respectively, TGA, 1393nt; TGA, 1396nt; TAA, 1397nt) in HBV genotype D. HepG2-NTCP cells^57^ were maintained in Dulbecco’s Modified Eagles Medium (DMEM) supplemented with 10% fetal bovine serum (FBS), 2 mM L-glutamine, 1 mM Sodium Pyruvate, 50 IU penicillin/streptomycin mL^−1^ and non-essential amino acids (Life Technologies, UK). Antibodies specific for HIF-1α were purchased from BD Biosciences (610959), anti-HIF-2α was purchased from Novus (NB100-132), and anti-CAIX was provided by the Harris laboratory (University of Oxford). HIF PHD inhibitor FG4592 was purchased from Cambridge Biosciences, UK. Cells were incubated under hypoxia in an atmosphere-regulated chamber with 1% O2: 5% CO2: balance N2 (Invivo 400, Baker-Ruskinn Technologies). The Ad-HBx and Ad-Ova express the HBV genotype D HBx gene and chicken ovalbumin gene under control of the Transthyretin (TTR) promoter. Promoter and insert were inserted into the E1 region of adenovirus (Ad5ΔE1/E3) backbone plasmid pAd/PL-DEST through Gateway recombination following the manufacturer’s instructions (Gateway System; Invitrogen, Karlsruhe, Germany). Adeno virus stocks were titrated using the cytopathic effect in HepG2 cells as previously described^58^.

### HBV genesis and infection

HBV was purified from a HepAD38 producer line as previously reported^57^. Briefly, virus was purified using centrifugal filter devices (Centricon Plus-70 and Biomax 100.000, Millipore Corp., Bedford, MA) and stocks with a titre between 3×10^9^ and 3×10^10^ viral genome equivalents (vge) per mL stored at −80°C. HBV-X-virus was purified from a HepG2 based cell line containing a HBV 1.3x overlength integrated viral genome where both 5’ and 3’ HBx genes were knocked out by a point mutation that changes the eight amino acid to a stop codon (CAA-to-TAA) as previously described^44^. HepG2-NTCP cells were treated with 2.5% dimethyl sulphoxide (DMSO) for 3 days and inoculated with HBV at an MOI of 200 in the presence of 4% polyethylene glycol 8000. After 18-20h the inoculum was removed by washing with PBS and the cells cultured in the presence of 2.5% DMSO. Secreted HBe and HBs antigen were quantified by ELISA (Autobio, China).

### PCR quantification of HBV RNA and HIF gene transcripts

Total cellular RNA was extracted using an RNeasy mini kit (Qiagen) following the manufacturer’s instructions and samples treated with RNase-Free DNaseI (14 Kunitz units/rxn, Qiagen) for 30 minutes at room temperature. RNA concentration was measured by NanoDrop 1000 spectrophotometer (Thermo Scientific) and cDNA synthesized with 0.25-1μg of RNA in a 20μL total reaction volume using a random hexamer/oligo dT strand synthesis kit in accordance with the manufacturer’s instructions (10 minutes at 25°C; 15 minutes at 42°C; 15 minutes at 48°C; SensiFast, Bioline). PCR amplification of HBV RNAs were performed using primers as previously described^43^ using a SYBR green real-time PCR protocol (qPCRBIO SyGreen, PCR Biosystems) in a Lightcycler 96™ instrument (Roche). The amplification conditions were: 95°C for 2 minutes (enzyme activation), followed by 45 cycles of amplification (95°C for 5 seconds; 60°C for 30 seconds). HIF target genes were amplified using TaqMan^®^ Gene Expression assays (CAIX [Hs00154208_m1]; VEGFA [Hs00900055_m1]; BNIP3 [Hs00969291_m1] and GLUT1 [Hs00892681_m1]) (Thermo Fisher) and amplified using a Taqman real-time PCR protocol (qPCRBIO probe, PCR Biosystems) using the same conditions as listed above.

### HBV transgenic mice and siRNA delivery

Animal experiments were conducted in accordance with the German regulations of the Society for Laboratory Animal Science (GV-SOLAS) and the European Health Law of the Federation of Laboratory Animal Science Associations (FELASA). Experiments were approved by the local Animal Care and Use Committee of Upper Bavaria and followed the 3R rules. Mice were kept in a specific-pathogen-free facility under appropriate biosafety level following institutional guidelines. HBVtg mice (strain HBV1.3.32)^59,60^ carrying a 1.3-fold overlength HBV genome (genotype D) on a C57BL/6J background and both male and female mice between 12-15 weeks were used. The HBV specific siRNA (siHBV) was designed to silence all HBV transcripts by targeting the 3’region of the HBV genome and the control siRNA (siCtrl) does not target any viral or known host transcripts. siRNAs were complexed with Invivofectamine 3.0 reagent (ThermoFisher Scientific) before injecting 1 μg/g body weight into the tail vain. HBeAg was quantified from mouse sera after dilution with the Architect HBsAg Manual Diluent using the quantitative HBeAg Reagent Kit (Ref: 6C32-27) with HBeAg Quantitative Calibrators (Ref.: 7P24-01) on an Architect TM platform (Abbott Laboratories, Wiesbaden, Germany). Immediately after sacrificing the mice and preparation of the liver, an approximately 0.4 mm thick and 1-1.5 cm long peace of liver was placed in 500μL RNAlater. After storage for 24h at 4°C (to allow RNA later to penetrate tissue) the tissue was transferred to −20°C and stored until RNA preparation. RNA was prepared using the RNeasy Mini kit (Qiagen), where an approximate 20mg piece of frozen liver was placed in a 2 mL micro-centrifuge tube pre-cooled on dry ice. After adding 600μL of Buffer RLT, tissue was homogenized using the TissueLyser LT (Qiagen) for 5 min at 50 Hz. Total RNA was extracted following the protocol of the RNeasy mini kit.

### HBV infected human chimeric mice and RNA-sequencing

Mock and HBV infected mice were sacrificed and livers harvested for RNA isolation and RNA-sequencing at the Beijing Genomics Institute (BGI, Hong Kong). RNA purity was assessed with a NanoDrop 2000 spectrophotometer (Thermo Fisher Scientific) and integrity determined using a 2100 Bioanalyzer Instrument (Agilent Technologies). Sequencing was performed on a BGISEQ-500 (Beijing Genomics Institute, Hong Kong) employing the PE100 mode to produce raw paired-end reads of 100 bp and SOAPnuke (v1.5.2) software to filter out non-human sequencing reads, as previously reported^50,61,62^. Clean reads (FASTQ files) were uploaded to Partek Flow (version 8.0, build 8.0.19.1125; Partek Inc., St. Louis, MO, USA), quality-controlled, and aligned to the human genome (hg38) with STAR - 2.6.1d aligner software. Genes were quantified using the transcript model Ensembl Transcripts release 91 and differential expression determined with DESeq2 (3.5). Microarray analysis was performed with Partek Genomics Suite (v6.6) as previously described^63^. Scatter dot plots of fold change values were plotted with Graphpad Prism version 8. RNA-seq data are deposited in the GEO archive at NCBI, with the accession number SE145835 and entitled: Transcriptional profiling of hepatocytes isolated from chronically HBV-infected human liver chimeric mice.

### Bioinformatic analyses

To determine whether HBV infection induces hypoxia-responsive genes, we interrogated the mRNA expression patterns of the liver chimeric mice RNA-Seq dataset for i) the top 80 hypoxia-induced genes as identified in HepG2 hepatic cells ^42^. In-house datasets were compared with RNA-seq^48^ from HBV-infected primary human hepatocytes. Data were retrieved from GEO (accessions: GSE120886, GSE93153, GSE118295). For consistency, all datasets were re-analysed with the same Partek Flow bioinformatic pipeline. Microarray analysis was performed with Partek Genomics Suite (v6.6) as previously described^63^ and data presented using Graphpad Prism 8.

#### Statistical Analyses

All analyses were performed using Prism 8 (GraphPad, La Jolla, CA). Data are shown as means ± SD, probabilities are indicated by * = p< 0.05, ** = p< 0.01, *** = p< 0.001 or **** = p<0.0001, with Bonferroni corrections for multiple testing when appropriate.

## ACKNOWLEDGEMENTS

We thank Peter Ratcliffe (University of Oxford) for his support and advice throughout this project and Peter Balfe for critical reading of the manuscript. Research in the McKeating laboratory is funded by Wellcome Trust IA 200838/Z/16/Z and MRC project grant MR/R022011/1. Collaborative research in the Protzer and McKeating laboratories was funded by EU Horizon 2020 program through the Hep-CAR consortium and the Institute for Advanced Study with the support of the Technische Universität München via the German Excellence Initiative and EU 7^th^ Framework Programme under grant agreement number 291763. TM was supported by a clinical leave stipend by the Else-Kröner Forschungskolleg “Microbial triggers as cause for disease” of Klinikum rechts der Isar, Technische Universität München. Research in the Klenerman lab is funded by Wellcome grant WT 109965MA and NIHR senior Fellowship. Research in Dorner laboratory was funded by an ERC grant (ERC-StG-2015-637304) and Wellcome Trust New Investigator award (104771/Z/14/Z).

## AUTHOR CONTRIBUTION STATEMENT

PJL designed and conducted experiments and co-wrote the manuscript; JMH designed and conducted experiments and co-wrote the manuscript; EM analysed data; VD conducted experiments; TM designed and conducted experiments and co-wrote the manuscript; PACW conducted experiments; AM analysed data; AM O-P conducted experiments; M vd K provided reagents, JW provided reagents; DD provided reagents; MD designed experiments; PK provided expertise; UP provided reagents; SG analyzed data and co-wrote the manuscript and JAM designed the study and co-wrote the manuscript.

## ADDITIONAL INFORMATION

RNA-seq data from HBV infected mice are deposited in the GEO archive at NCBI, with the accession number SE145835 and entitled: Transcriptional profiling of hepatocytes isolated from chronically HBV-infected human liver chimeric mice. Our in-house data was compared with RNA-seq^48^ from HBV-infected primary human hepatocytes and data retrieved from GEO (accessions: GSE120886, GSE93153, GSE118295).

---

**Supplementary Table 1.**
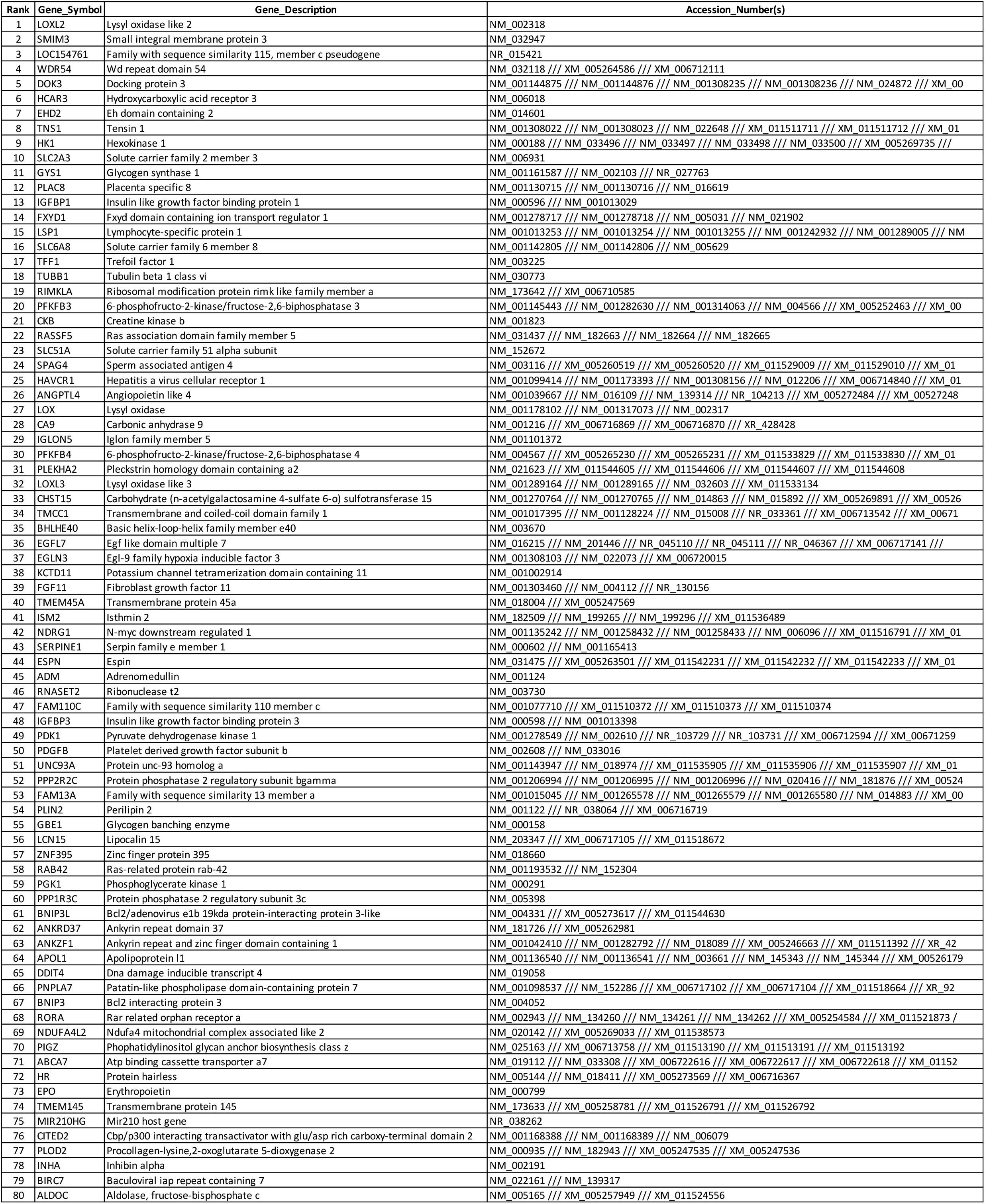
80 Hypoxia signature genes

